# Fascicles Split or Merge Every ~560 Microns Within the Human Cervical Vagus Nerve

**DOI:** 10.1101/2021.11.09.467343

**Authors:** Aniruddha R. Upadhye, Chaitanya Kolluru, Lindsey Druschel, Luna Al Lababidi, Sami S. Ahmad, Dhariyat M. Menendez, Ozge N. Buyukcelik, Megan L. Settell, Stephan L. Blanz, Michael W. Jenkins, David L. Wilson, Jing Zhang, Curtis Tatsuoka, Warren M. Grill, Nicole A. Pelot, Kip A. Ludwig, Kenneth J. Gustafson, Andrew J. Shoffstall

## Abstract

Vagus nerve stimulation (VNS) is FDA approved for stroke rehabilitation, epilepsy, and depression; however, the vagus functional anatomy underlying the implant is poorly understood. We used microCT to quantify fascicular structure and neuroanatomy within human cervical vagus nerves. Fascicles split or merged every ~560 μm (17.8 ± 6.1 events/cm). The high degree of fascicular splitting and merging in humans may explain the clinical heterogeneity in patient responses.

## 2 Main

Electrical stimulation of the cervical vagus nerve (cVN) using implanted electrodes, more commonly known as cervical vagus nerve stimulation (cVNS), is an existing clinical therapy with an estimated global market size of over $500 million dollars in 2018. This market is projected to expand at a compound annual growth rate of 11.4% to a size of nearly 1.2 billion dollars by 2026.^1^ Implanted vagus nerve stimulators are currently approved by the Food and Drug Administration (FDA) to treat epilepsy, depression, obesity and for stroke rehabilitation^2–5^, and are in clinical trials to treat diverse conditions including heart failure, diabetes, and rheumatoid arthritis.^6–8^

The vagus nerve at the cervical/neck level is an attractive target for neuromodulation therapies as it is easily identifiable under ultrasound and can be accessed with a well-established and relatively simple surgical procedure.^9^ In humans, the cervical vagus consists of over 100,000 fibers; these include efferent fibers originating from the brainstem that innervate multiple visceral organs, including the lungs, heart, diaphragm, liver, and intestines, and their sensory fibers returning to the brainstem, which ultimately influence noradrenergic, serotonergic, and cholinergic inputs to the cortex.^9–11^ As such, intervening at the cervical vagus presents the opportunity to modify function both within the brain and the majority of organs within the viscera.^12–21^

Several recent studies in animal models have suggested that smaller, multi-contact electrodes may more selectively stimulate specific portions of the cervical vagus to take advantage of underlying functional organization to better isolate intended activation of therapeutic fibers from unwanted activation of off-target fibers.^22,23^ The activation of low-threshold, large-diameter motor efferent fibers of the vagus that innervate the deep muscle of the necks putatively drives the most common side effects, causing cough, throat pain, voice alteration, and dyspnea reported in up to 66% of patients.^24–29^ In a study of human patients implanted to treat heart failure, desired heart rate responses were achieved in only 13 of 106 measurements taken at the 6-and 12-month end points, with stimulation thresholds predominantly limited by side effects attributable to concurrent activation of the neck muscles.^24^

The vagus nerve is known to have distinct functional organization at specific points along its path connecting the brainstem to the visceral organs.^30,31^ Motor efferents responsible for deep neck muscle activation originate within the nucleus ambiguus in the medulla oblongata and eventually coalesce into the pharyngeal, superior laryngeal, and recurrent laryngeal branch, which innervate the pharyngeal, cricothyroid muscle, and cricoarytenoid muscles, respectively. Parasympathetic efferents originate from the dorsal motor nucleus of the vagus within the medulla oblongata and travel down the cervical vagus and eventually join vagal branches leading to and from the visceral organs. In contrast, sensory afferents leading from the visceral organs follow these same branches back to the main trunk that eventually becomes the cervical vagus.

While much is known about the proximal/distal connectivity of the vagus nerve, it is unknown if the human vagus at the cervical level has well-maintained functional organization, or lack thereof, that may account for the high degree of heterogeneous results across patients clinically. Seminal studies by Sunderland have previously demonstrated that although the fascicles of major peripheral nerves divide and unite to form fascicular plexuses, there is substantial uniformity of fascicular arrangement of major nerves in the extremities.^32,33^ For example, the palmar cutaneous and motor branches of the median nerve can be dissected proximally for several centimeters without significant cross branching.^32,33^ Prior studies in human cadavers have focused on sparse sampling of the cervical vagus and subsequent 2D sectioning, which has yielded highly variable results with respect to number of fascicles from study to study with little information about the underlying functional somatotopy relevant to VNS.^34–37^

In this study, we collected 8 mid-cervical VNs from 5 human cadavers; each nerve was 5 cm long, and we focused our quantitative analyses on the middle 1 cm where the clinical VNS cuff would be surgically placed.^38^ We stained the nerves with osmium tetroxide, and we imaged the nerves’ morphology in three dimensions using microCT. We visualized and quantified the merging and splitting of fascicles along the 1 cm window (**Figures 1, 2**). Merging and splitting events were detected manually by an impartial observer (**Figure 1 A, C**), noting delineation by perineurium boundaries (**Figure 1 B**). We measured the distance over which the events occurred; merges spanned 430 ± 117 (μm ± SD, n = 70) and splits spanned 461 ± 108 (n = 72) (**Figure 1 D**).

**Figure 1:**
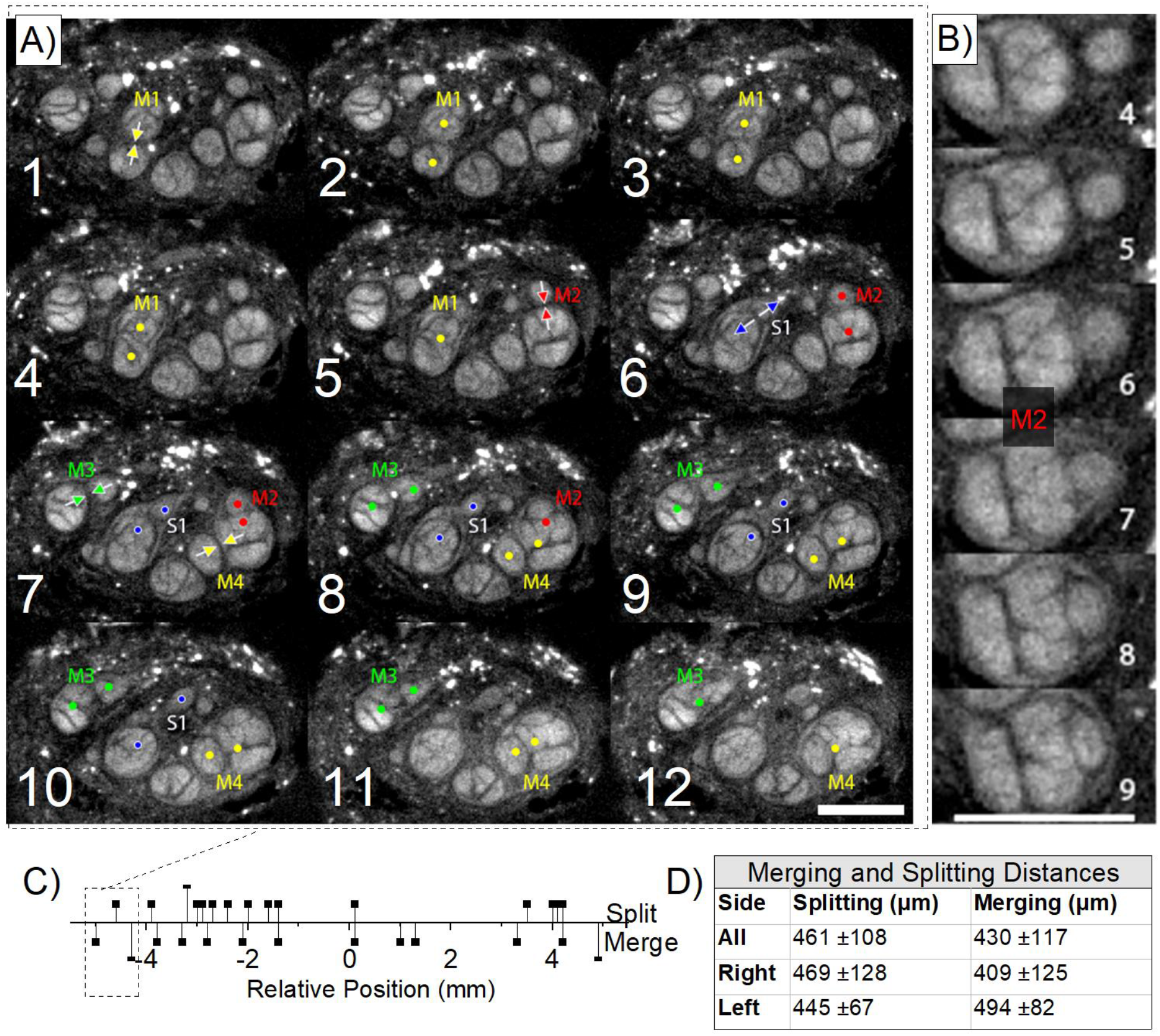
Representative example of splitting and merging of fascicles along the rostral-to-caudal direction within a 1. 1 mm length of the human cVN (Specimen “2R”) imaged with microCT. A) The initiation of merging “M” and splitting “S” events are annotated with arrows: 4 merges (M1-M4) and 1 split (S1). Frames are read from left-to-right, top-to-bottom, as if reading text. Frame-to-frame spacing is 100 μm (12 frames = 1.1 mm total longitudinal span). Transverse-plane scale bar shown in bottom right of the figure is 500 μm. B) Example merging event “M2”, spanning 6 frames (500 μm). C) A representative line graph depicting event frequency (Split-positive, Merge-negative) along the middle 1 cm length of nerve. D) Table of mean distances (mean ± SD) over which split and merge events (n=72 and n= 70, respectively) occur for all 8 VNs, sampled from either from the right or left side of the neck (middle 1 cm).

**Figure 2:**
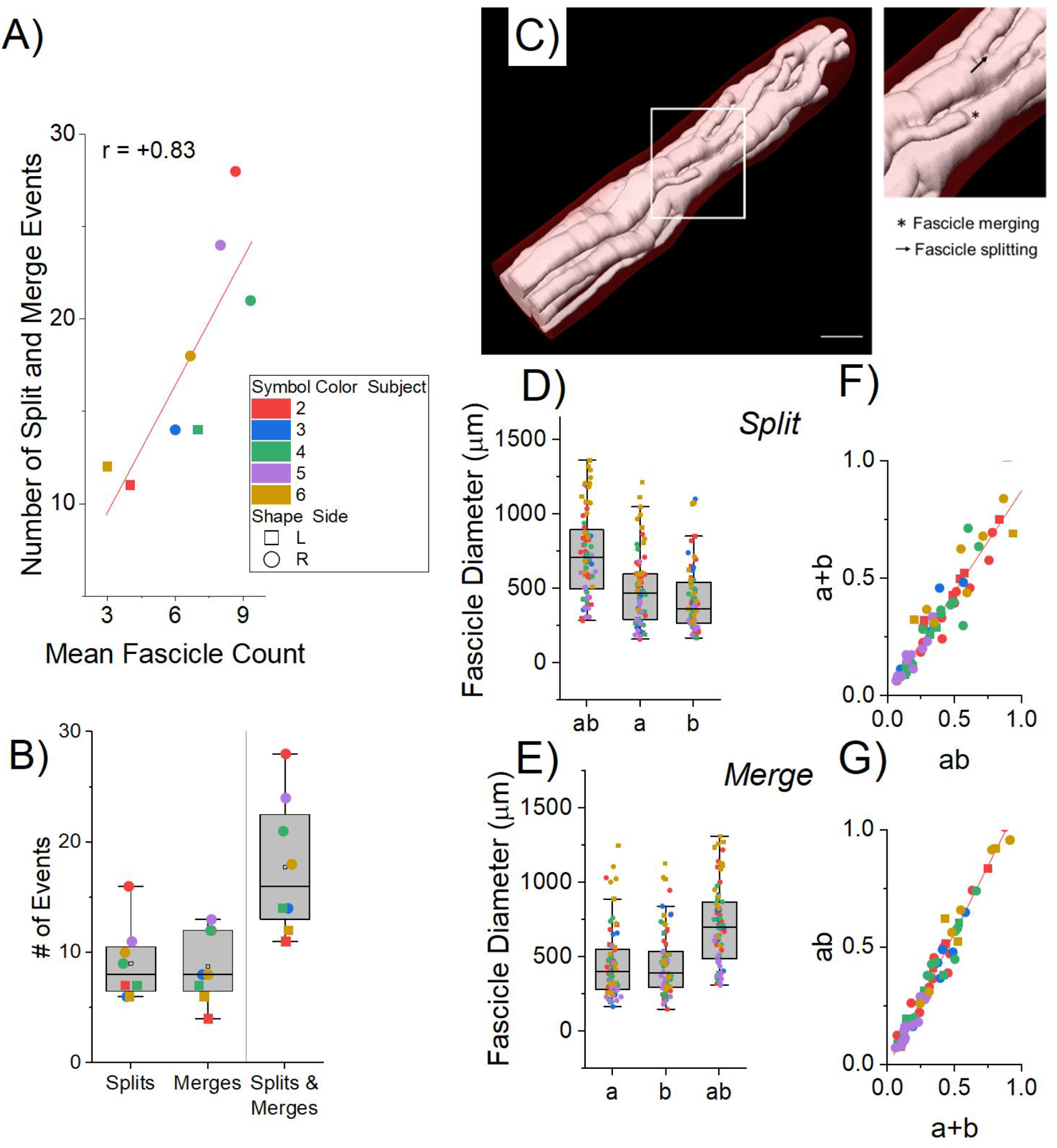
Graphical representation of fascicular dynamics within the central 1 cm of the surgical window for VNS implantation across 8 nerves. The quantification of these events was possible due to the high resolution along the longitudinal axis of the microCT dataset. A) Correlation between the number of fascicles and the number of split/merge events along the 1 cm length of nerve: subject number (color-coded, 2 – 6), left (square), right (circle). B) Box plot showing the distribution of the number of split/merge events across all samples. C) 3D visualization of a representative 1 cm window within the cVN (Specimen “4R”). D, E) Box plot showing the distribution of the diameters of parent fascicles and children (a, b or a+b, respectively) for all merge and split events. F) Association plot of splitting fascicular summed areas of the children (a+b, y-axis) with the areas of the parent (ab, x-axis), mixed model slope β=0.87, p<0.001. G) Association plot of merging fascicular areas of the parent (ab, y-axis) with the summed areas of the children (a+b, x-axis), mixed model slope β=1.14, p<0.001. Note that summed areas of the children are consistently less than the area of the parent fascicle.

Over the middle 1 cm of all 8 nerves, there were 17.8 ± 6.1 merging and splitting events (**Figure 2 B, C**), meaning that on average, each fascicle split or merged every ~560 μm. This number of events is much larger than expected from prior studies using histological techniques.^34,35,37^ For the standard clinical VNS cuff electrode (LivaNova, London, UK) and a nerve with ~6.6 fascicles (the mean value in our study), one would expect to observe ~14.2 split or merge events over the 8 mm between the centers of the bipolar contact pair. These rapid shifts in fascicular organization would be challenging to observe using standard histological or electron microscopy methods—typically using a single transverse cross section per nerve—and thus, prior studies on vagal morphology have not quantified this phenomenon.^35,37^

Merging and splitting events increased proportionally with the number of fascicles: more fascicles provided more opportunity for split/merge events (**Figure 2 A**, β = 1.76, *p* = 0.032). We used a two-level linear mixed model considering subject and spatial correlation between samples to evaluate for association. This degree of fascicular reorganization has substantial implications for VNS due to changing perineurium boundaries, which dramatically influences the distribution of the electric field.^39^ The locations of fibers—and therefore proximity of fibers to the electrode contacts—also directly influences activation thresholds. Fascicles of a wide range of diameters participated in splitting and merging events; reorganization was not limited to a sub-population of small or large diameter fascicles (**Figure 2 D, E**).

Additionally, the cross-sectional areas of parent (“ab”) and summed children (“a” + “b”) fascicles before and after merging or splitting events (**Figure 2 F, G)** were calculated and compared (i.e., “ab” vs “a + b”). The parent areas were consistently larger than the sum of the children areas (β = 0.87, *p* <0.001 and β = 1.14, *p* <0.001, for splitting and merging, respectively, where β refers to the slope of the mixed model).

Using the microCT images, we generated a 3D model (**Figure 3 A**) and quantified the fascicular morphology: number of fascicles, effective circular diameter, and cross-sectional area (**Figure 3 B-G**). Statistically, there was a net increase in mean fascicle diameter (*p*=0.0139) in the cranial to caudal direction **(Figure 3D, E)** with negligible change in overall fascicular area (*p*=0.8399, **Figure 3 F, G**), suggesting a consolidation of the fascicles toward the inferior end of the cervical region. However, the large subject-to-subject variability is the overwhelming takeaway from the data (**Figure 3**). We did not observe any branches, although branches may occur in this region in some individuals.^34^ While there was a trend toward a concomitant decrease in fascicle count with longitudinal distance (**Figure 3B, C)**, the result was statistically insignificant using a mixed-effects regression model (*p*=0.1672, data not shown), likely owing to low sample number and the substantial variation between subjects for all three morphological metrics.

**Figure 3:**
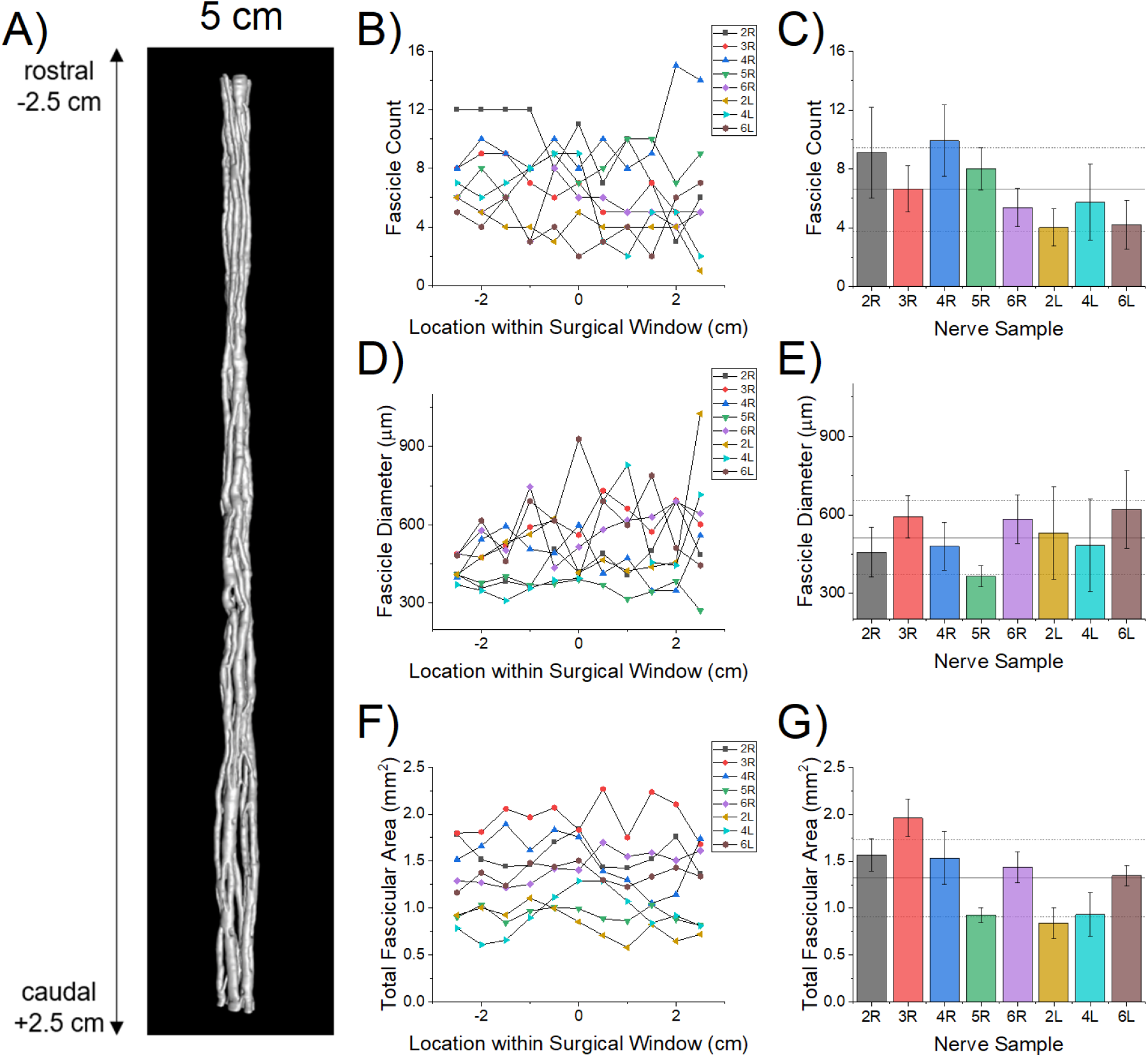
Fascicle morphometry assessment within the central 5 cm of the human cVN (Specimen “4R”). A) Representative 3D visualization of segmented microCT images. B, D, E) Fascicle count, diameter, and area at 0.5 cm increments along the 5 cm surgical window for each sample, where x = –2.5 cm is the rostral end and x = +2.5 cm is the caudal end. We also averaged the data across the surgical window for each sample (C, E, G). Bars represent the mean ± SD across the sampled regions of the surgical window. Black horizontal lines represent the mean ± SD across all nerve samples.

MicroCT enables unique three-dimensional visualization and quantification of vagal fascicular morphology over long lengths of nerve, enabling new insights into the spatial organization of the nerve that are essential for the design and analysis of effective and selective electrical stimulation therapies to treat diseases. MicroCT has been used extensively in orthopedic studies and other fields, but the imaging technique has only recently been applied to neural tissues. For example, one study reported a protocol for staining rat sciatic and pig vagus nerves, optimization of computational methods for high-resolution three-dimensional images of nerve fascicles, and development of image analysis techniques to facilitate segmentation and tracing of the fascicles.^40^ The fascicle morphology measurements obtained from our microCT data were similar to those obtained by other groups.^37^ Here, we demonstrated the unique value of microCT to quantify fascicular splitting and merging of the human cVN.

Given the magnitude of fascicular reorganization demonstrated by our data, current VNS cuff designs are not optimized to provide spatial selectivity. The current clinical standard involves surgical implantation of a cuff electrode that wraps helically around the entire nerve trunk, with bipolar contacts spanning ~270°, separated by 8 mm center-to-center. For a representative nerve from our study, this 8 mm span would traverse over a dozen fascicle splitting and merging events (min = 9.6, max = 22.4 events, from our limited size dataset). Further, the fascicular reorganization varies substantially between individuals. Given this intra-and inter-individual morphological heterogeneity of fascicles, these electrode designs are unlikely to allow selective activation of spatially localized target fibers within the cVN.

Computational modeling of the vagus nerve can be used to guide the engineering and design of neural stimulating devices^41^; the basis for these models requires anatomically accurate features that reflect the diversity observed across multiple human subjects. Currently, computational modeling of VNS relies on longitudinal extrusion of segmented histological cross sections or simplified mock morphologies, which do not represent precise fascicle boundaries or longitudinal spatial variation. ^42–44 45 46,47^ Autonomic stimulation therapies will be advanced by *a priori* personalized surgical planning, device design, and device programming for autonomic stimulation therapies informed by computational models as used in other neural stimulation treatments.^48^ However, to make personalized decisions and improve the accuracy of the computational predictions, better *in vivo* imaging modalities are needed to visualize and map the fascicular morphology with higher precision and resolution in both the transverse and the longitudinal planes.^49^

The fascicular anatomy of vagus nerve is highly complex and dynamic. Mapping nerves using microCT is an effective technique to visualize and quantify fascicle reorganization. We measured a mean of 17.8 split-or-merge events along 1 cm of the cervical vagus nerve (n=8 samples), implying that there would be ~14 events along the bipolar electrode of current clinical VNS devices. The analysis of fascicle dynamics within the human VN provides a unique perspective into the morphology of the VN and suggests that morphology may have implications on VNS efficacy. Specifically, this analysis provides the foundation for building computational models to analyze and design therapies with improved selectivity reducing off target effects which can greatly improve patient’s quality of life. Such therapies could lead to an overall improvement in clinical outcomes.

## 3 Methods

### 3.1 Tissue Acquisition and Dissection

We collected 8 mid-cervical vagus nerve samples from 5 formaldehyde fixed cadavers (3 left nerves, 5 right nerves), secondary to use in medical school cadaver lab training. Since all the specimens were harvested from de-identified donor sources, and no protected personal health information collected, a letter of IRB exemption (non-human-subjects determination) was sought and approved by the Case Western Reserve University Institutional Review Board.

Cadavers were already disarticulated prior to our dissection; we performed gross and fine dissection with standard tools to isolate the vagus nerve from surrounding tissues. We made a rostral cut directly beneath the skull (jugular foramen) approximately at the nodose ganglion. The caudal/distal cut was made at the level of clavicle. The harvested nerves were stored in 4% formalin solution until ready for staining. The VNS cuff electrode is clinically placed midway between the clavicle and the mastoid process, and the surgical incision is 3-4 cm long^38^; we therefore collected 5 cm of length for each nerve, centered around the approximate location of VNS cuff placement, which we refer to as the “surgical window” throughout the paper.

### 3.2 Sample Preparation: Osmium Staining & Paraffin Embedding

The vagus nerves were washed three times with 1X phosphate buffered saline (PBS), letting the sample shake on an orbital shaker for five minutes after each wash. Osmium tetroxide (1% v/v) was prepared with deionized water, and the nerves were left fully submerged in this solution for three days. The samples were then dehydrated with 70% and 95% ethanol with a deionized water solvent. The dehydration included two quick rinses of the samples with 70% ethanol followed by a full wash and 30 minutes on the orbital shaker. This process was repeated twice with 70% ethanol, then three additional times with 95% ethanol. The nerves were stored in 70% ethanol for up to one week prior to embedding in paraffin.

The nerve samples were embedded in paraffin, mounted on a 3D printed plastic mold that fit the nerve. At the base of the mold, there were grooves every 5 mm, and these grooves were painted with a marking solution doped with barium sulfate to enhance sample navigation under X-ray.

### 3.3 MicroCT and Image Sub-Volume Reconstruction

For the imaging studies, we utilized a Quantum GX2 microCT Imaging System (Perkin Elmer, Waltham, MA, USA). The embedded nerve was placed in a 36 mm bed. The microCT scanner was warmed up as recommended by the manufacturer, and the nerve was scanned and reconstructed at 36 mm field of view (FOV). The resultant image block was 72 μm in voxel resolution (isotropic). Each scan spanned 1.8 cm of nerve length, with 0.3 cm overlap (i.e., 16.67%) between adjacent blocks to serve image reconstruction.

Post-hoc sub-block reconstruction was performed with Rigaku software provided by Perkin Elmer. Each sub-block reconstruction was a 5.12 × 5.12 × 5.12 mm^3^ cube and each adjacent sub-blocks overlapped by 0.1 mm (20% overlap); the resolution of final reconstruction was 10 μm voxel size (isotropic). Images were exported as DICOM files for further processing. After down-sampling frames along the longitudinal axis by 10-fold, blocks were co-registered and stitched using ImageJ (FIJI, Version 2.1.0/1.53c).^50^ The final image dataset consisted of pairwise stitched, evenly spaced (100 μm inter-frame spacing) TIFF images. 3D visualizations were generated by Simpleware™ ScanIP software (Synopsys, Mountain View, CA).

### 3.4 Fascicle Morphometric Analysis

VN samples were analyzed using ImageJ (FIJI, Version 2.1.0/1.53c) to select, outline, and measure individual fascicles, using the elliptical selection tool. Fascicle boundaries were manually estimated based on visual inspection. For morphometric analysis, the operators evaluated fascicle parameters at 0.5 cm intervals along the length of the 5 cm cervical window for each nerve. While manual outlining potentially introduces subjective differences between operators, the magnitude of these differences was deemed negligible based on prior inter-operator analyses. Image scaling was set according to the microCT manufacturer provided calibration factor: 1 pixel = 10 μm, 1.0-pixel aspect ratio. Area, minimum & maximum gray intensity values, shape descriptors, mean intensity value, centroid coordinates, and ellipse-fit measurements (including major and minor axes, and effective diameter – the average of major and minor axis) were calculated.

### 3.5 Merging and Splitting Analysis

The splitting and merging analyses were conducted for the central 1 cm of the cervical vagus nerve, within the 5 cm of the surgical window that we defined in this paper. The frames in this region were isolated and loaded as an image sequence on ImageJ and analyzed from the rostral end to caudal end. All split/merge analyses were conducted manually.

#### 3.5.1 Defining an Event

During our analysis, we defined the start and completion of a split or merge event based on the fascicle boundaries. We characterized an event as a start of a split when a parent fascicle, coined “ab”, appeared to create a bud or partition within the center of the otherwise consistently shaded fascicle (e.g., **Figure 1B**). The event was marked as complete when parent fascicle “ab” completely formed independent circular/ellipsoidal independent children fascicles “a” and “b” with their own perineurium sheath around the fascicles. In most cases, the perineurium sheathe is well defined and visible within the microCT. In some cases, the perineurium is inferred when there is separation of two circular/ellipsoidal geometries. Conversely, we characterized an event as a merge when fascicle “a” merged into another fascicle “b”, resulting in a combined fascicle “ab”, applying the same logic as described above. When multiple events occurred simultaneously (e.g., one fascicle splitting into three fascicles), we considered it as two different splitting events. We did not observe any event where three fascicles merged to become one fascicle in the exact same frame.

#### 3.5.2 Measurements and Analysis

To measure the distance over which the event was taking place, the starting and the ending frames were recorded. With the total number of frames, we calculate the distance over which the event takes place. Using ImageJ, the fascicles were measured at the starting and the ending frames (as mentioned in the morphometric analysis section).

We recorded the number of splitting and merging events across the central 1 cm of each sample and calculated the average number of events across n = 8 samples. We counted the number of fascicles in the first, middle, and last frames of the 1 cm window and calculated the mean fascicle count in the sample. We then determined the number of events/fascicle/cm using the values calculated as mentioned previously.

### 3.6 Statistics

Our primary quantitative metric was focused on fascicle splitting and merging events across our human cadaver nerve specimens (n = 8). Descriptive statistics presented in the text include mean and standard deviations unless otherwise denoted. Box plots presented in **Figure 2** display individual data points (colored according to the associated legends), median values (horizontal center line), mean values (black box), interquartile range (upper and lower box edge), and outliers (whiskers). Bar plots presented in **Figure 3** display mean values (bar height) and standard deviation (error bars), with horizontal lines in the background representing the whole sample mean and standard deviations.

For all statistical tests described below, Two-sided Type I error = 0.05 was adopted. Analysis was performed using R version 4.0.2.

Specifically, we were interested in evaluating the relationship between the number of fascicles contained within nerve specimens and the number of splitting or merging events observed (**Figure 2 A**). The association between the average number of fascicles at the surgical window and the number of events along the window was investigated with a two-level linear mixed model with subject and (left or right) side-level random intercepts.

We were also interested in evaluating the conservation of fascicular area before-and-after splitting and merging events (**Figure 2 F, G**). In order to study the association between fascicular area of the parent (ab) and summed areas of the children (a+b), we adopted a three-level hierarchical linear mixed model with subject-level and side-level random intercept with exponential spatial correlation structure for same side windows.

Similar 3-level models, as described above, were respectively used to explore the spatial trend of outcomes along the surgical window (rostral-to-caudal) for fascicular area, mean diameter, and fascicle count (results shared in text).

### 3.7 Methodological Limitations

As with standard histological processes, the staining and fixative reagents can cause dehydration and shrinkage to tissues. Per prior publications, we anticipate shrinkage could contribute up to 20% reduction in apparent diameters. However, we did not directly estimate this in our study, and therefore did not apply any correction factors in our dataset. Further, we sampled nerves from 5 cadavers, but due to the source of cadaver donation, we were unable to acquire any demographics. This study can be expanded in the future to greater population sample size to estimate population variability drive by demographic differences.

## 4 Acknowledgements

The authors would like to thank Rebecca Enterline and Andrew Crofton for their contributions in sample acquisition and handling. We would like to thank Matt Schiefer for his role in the acquisition of equipment necessary for the execution of our studies. We would also like to recognize William Woodfint for his contributions to data review.

This work has been supported by NIH SPARC Program 1OT2OD025340, US Dept. of Veterans Affairs 1IS1BX004384, the Cleveland VA APT Center, and Case Western Reserve University. The opinions expressed in this article are the author’s own and do not reflect the view of the National Institutes of Health, the Department of Health and Human Services, or the United States government.

